# Regulatory FOXP3+ T cells in uterine sarcomas are associated with favorable prognosis, low extracellular matrix expression and reduced YAP activation

**DOI:** 10.1101/2021.03.26.437169

**Authors:** Okan Gultekin, Jordi Gonzalez-Molina, Elin Hardell, Lidia Moyano-Galceran, Nicholas Mitsios, Jan Mulder, Georgia Kokaraki, Anders Isaksson, Dhifaf Sarhan, Kaisa Lehti, Joseph W. Carlson

## Abstract

**Purpose:** Uterine sarcomas are rare but deadly malignancies without effective treatment. The goal of this study was to characterize and identify potential mechanisms underlying observed variations in the immune microenvironment of different sarcoma subtypes, using integrated clinicopathological and molecular methods.

**Experimental design:** Fifty-eight cases of uterine sarcoma with full clinicopathological annotation were analyzed for their immune landscape in the tumor microenvironment, gene, and protein expression. Cases included leiomyosarcoma (LMS; n=13), low-grade endometrial stromal sarcoma (ESS; n=16), undifferentiated uterine sarcoma (UUS; n=26), and YWHAE-FAM22 translocation-bearing ESS (YFAM; n=3). Image analysis was used to quantify immune cells and immune regulatory proteins. Gene ontology and network enrichment analysis of matching transcriptomic data was used to relate over- and under expressed genes to pathways and further to the immune phenotype and clinicopathological findings.

**Results:** Immune cell characterization revealed overall prevalence of regulatory T cells and the pro-tumor M2-like macrophages. Cytotoxic T cells were only found in ESS and UUS tumors. Expression of immune regulatory proteins was heterogeneous, with PD-L1 being undetectable. Hierarchical clustering of patients showed four immune signatures independent of tumor type, where infiltration of non-exhausted FOXP3^+^ cells and M1-like macrophages were associated with greater overall survival. High CD8^+^/FOXP3^+^ ratio in UUS and ESS was associated with poor survival and upregulation of extracellular matrix (ECM)-related genes and proteins and YAP nuclear localization.

**Conclusions:** Uterine sarcomas present distinct immune signatures with prognostic value, independent of tumor type. This study suggests that the ECM is a potential regulator of the immune microenvironment in uterine sarcomas.

## Introduction

Uterine sarcomas are a heterogeneous group of rare neoplasms comprising 3-4 % of all uterine malignancies (1). Despite their rareness, they are responsible for considerable mortality and morbidity by frequently recurring and metastasizing distantly (2). Uterine sarcomas are classified into various histopathological subtypes, and treatment is subtype specific (3). These subtypes include the most frequently diagnosed, uterine leiomyosarcoma (LMS), and rarer subtypes such as low-grade and high-grade endometrial stromal sarcoma (LG-ESS and YFAM), adenosarcoma and undifferentiated uterine sarcoma (UUS) (1,2,4). Current treatments include surgery, chemotherapy, radiotherapy, and hormone therapy. However, their poor effectiveness highlights the urgent need for new therapies (2,5,6)

Immunotherapies are promising treatments that exploit the immune system to eliminate cancer cells. Immune evasion is essential for cancer cells to survive and is achieved by manipulating immune checkpoint pathways, normally used to maintain self-tolerance (7). Checkpoint inhibitors have shown promising results in the treatment of various tumor types. The efficiency of these treatments in uterine sarcomas remains low (8).

The success of immunotherapies relies on the ability to manipulate the tumor immune microenvironment (TIME) into an anti-tumorigenic state. Checkpoint inhibitor treatments can accomplish this, but their efficacy depends on the expression of targetable immune regulatory proteins (IRPs) and the cellular composition of the TIME. Among these IRPs, the PD-1/PD-L1 pathway, IDO1 and B7-H4 have sparked special interest (7,8)

The immune cell repertoire of the TIME is another determining factor for the efficacy of immunotherapies. Tumors rich in CD8+ cytotoxic T lymphocytes (CTLs), frontline defensive cells for targeting and killing tumor cells, often are associated with good prognosis and immunotherapy response (9). Moreover, targeting exhausted CTLs can enhance antitumor immunity (10). Other promising targetable components of the TIME include regulatory T cells (Treg), CD4+ T cells, and macrophages. The immune inhibitory function of Tregs is generally considered to have pro-tumorigenic effects (11,12). Similarly, anti-inflammatory or M2-like macrophages are associated with poor prognosis while pro-inflammatory or M1-like macrophages are a marker of good prognosis in various cancer types (13,14).

In this study we aimed to comprehensively characterize the TIME in uterine sarcomas in order to identify immune signatures with potential for current and future immunotherapies.

## Materials and Methods

### Patient cohort and central review

The retrospective cohort used in this study contains material from two collaborating centers, the Karolinska University Hospital and Skånes University Hospital. Ethical approval was obtained from the relevant authorities. All cases were reviewed centrally, and the cases of endometrial stromal sarcoma and UUS were analyzed with RT-PCR for detection of the YWHAE-FAM22 translocation. Details of the histopathological review have been described previously at Binzer-Panchal et al (15).

### Multiplex immunofluorescence

Tissue microarray (TMA) slides were air dried at room temperature, deparaffinized and then rehydrated in phosphate buffered saline (PBS) (P4417 Sigma-Aldrich, St. Louis, MO, USA) for 15 min. Primary antibodies (Supplementary Table 1) were diluted in 0.3% Triton X-100 (X100, Sigma-Aldrich, St. Louis, MO, USA) containing 0.1% NaN3 in PBS pH7.4 and applied to the slides for 16 h (4°C). Afterwards, sections were washed three times in Tris Buffered Saline solution, containing Tween^®^ 20 (0.05%) (TBS-Tween 20, T9039 Sigma-Aldrich, St. Louis, MO, USA) for 15 min each. The sections were then incubated in a Tris–NaCl-blocking buffer (TNB-buffer, FP1020, PerkinElmer, Waltham, MA, USA) for 30 min at room temperature. This was followed by the addition of the secondary fluorescent labeled antibody mix (Supplementary Table 1) diluted in TNB-buffer and sections were incubated for 90 min at room temperature. Subsequently, the sections were washed three times in 0.05% TBS-Tween 20 for 15 min, in the dark. The sections were then immersed in 70% EtOH for 5 min before being transferred to Sudan Black (1% solution, 70%EtOH) for 10 min and then rinsed in 70% EtOH for about a minute before mounting, using PVA/DABCO medium (ProLong^®^ Gold anti-fade with DAPI, P36931, Life Technologies, Thermo Fisher Scientific, Waltham, MA, USA).

Fluorescent images were obtained using a “VSlide” slide scanning microscope (MetaSystems, Alltlussheim, Germany). The system has a CoolCube 2 camera (12-bit grayscale), a 10x objective and filter sets for 4′,6-diamidino-2-phenylindole (DAPI) (EX350/50–EM470/40), Fluorescein isothiocyanate (FITC) (EX493/16−EM527/30), Cyanine (Cy) 3 (EX546/10−EM580/30), Cy3.5 (EX581/10−EM617/40), and Cy5 (EX630/20−647/long pass). First, the whole TMA was initially pre-scanned at 2.5x to generate the TMA scanning area. Tissue and focus depth were detected based on the DAPI signal. All tissue-covered areas were scanned using a 10x objective. Finally, the individual images were stitched together (VSlide) to generate a large image of the entire section. After scanning, the images (vsi-files) were extracted to high quality Iconforge Create Executable Library Data (ims)-files for further analysis using the software Qupath (Metasystems, Alltlussheim, Germany). In order to facilitate the analysis, the images were not downsampled to avoid losing any valuable biological information.

### Immunofluorescence quantification

Qupath (v.0.2.0-m2) (https://github.com/petebankhead/qupath) open source soft-ware (16), was used for single cell detection on 3 TMAs with IF staining for CD8, FOXP3, PDCD1, CD68, CD163, DAPI. TMA slides that hold 80 of the cases were dearrayed, and then detected.

### Unsupervised patient clustering

The average cell densities of two cores from each patient was used to create unsupervised patient clustering. The analysis was conducted in R v.4.0.3 (R Core Team, 2020) with heatmap.2 package (Galili, 2020).

### Gene expression analysis

The RNA expression arrays method has been previously described (15). Briefly, RNA from each sample was used to generate amplified and biotinylated sense-strand cDNA from the entire expressed genome according to the Sensation Plus FFPE Amplification and WT Labeling Kit (P/N 703089, Rev.4 Thermo Fisher Scientific Inc., Life Technologies). GeneChip ST Arrays (GeneChip Human Gene 2.1 ST Array Plate) were hybridized, washed, stained, and finally scanned with the GeneTitan Multichannel (MC) Instrument, according to the GeneTitan Instrument User Guide for Expression Array Plates (PN 702933, Thermo Fisher, Scientific Inc., Life Technologies).

The RNA raw data were normalized and compared using the free Affymetrix Expression Console Software provided by Thermo Fisher. Pathway analysis of differentially expressed genes was conducted by Metascape (17).

### Immunohistochemistry

The method for immunohistochemistry (IHC) staining of MMP14, collagen I, collagen IV and fibronectin was previously described (15). Briefly, TMA sections were deparaffinized and rehydrated. Antigen retrieval was performed using 10 mmol/L sodium citrate pH 6.

Endogenous peroxidase was quenched with 0.6% H2O2 for 10 minutes, 2 × 5 min PBS (for ImmPRESS kit) or with 0.03% H_2_O_2_ for 10 minutes, 1 minute H2O, 10 minutes PBS [for Tyramide Signal Amplification (TSA) kit].

The ImmPRESS method was used for MMP14 staining with anti-MT1-MMP (LEM) antibody (Supplementary Table 1). The TSA method was used for collagen I, collagen VI, and fibronectin stainings (antibody information in (Supplementary Table 1).

Finally, CD4 and YAP stainings (antibody information in Supplementary Table 1) were performed on tissue sections retrieved at the accredited clinical laboratory of the Department of Pathology, Karolinska University Hospital, Sweden. Staining was performed in the routine pathology laboratory by using an automated Ventana Benchmark Ultra system (Ventana Medical Systems, Tucson, AZ, USA).

### Statistical Analysis

Descriptive statistics were calculated and presented in tables. For the comparison of the density within groups and intragroup, non-parametric ANOVA (Kruskal-Wallis test) and one-way ANOVA testing were used, respectively. The overall survival (OS) probabilities were estimated and presented by Kaplan-Meier survival curves. The correlation between densities was obtained using Spearman’s rank correlation. All statistical tests were two-sided. P values less than 0.05 were considered statistically significant. Prism v8.0 software (GraphPad) and R programming were used for statistical analysis.

## Results

### Patient Cohort

Clinicopathologic characteristics are presented in Supplementary Table 2. A total of 58 malignant tumor cases were included, comprising 13 leiomyosarcomas (LMS), 16 endometrial stromal sarcomas (ESS), 26 undifferentiated uterine sarcomas (UUS), and 3 YWHAE-FAM22 endometrial stromal sarcomas (YFAM). In addition, there were 14 benign leiomyomata controls (LM). The mean follow-up time for LMS was 4.99 years (range 0.55-16.42), for ESS 10.88 years (range 2.93-20.67), for UUS 5.02 years (range 0.09-20.38), and for YFAM 11.16 years (range 2.61-18.79). For LMS 11/13 (84.6%) patients were deceased at last follow-up, for ESS 2/16 (12.5%), for UUS 22/26 (84.6%), and for YFAM 2/3 (66.7%).

### Multiplex immunofluorescence reveals T cell and macrophage heterogeneity across sarcoma subtypes

To characterize the TIME in uterine sarcomas, tumors were analyzed for T cell and macrophage infiltrates, and for the expression of immune regulatory molecules. Characterization of T cells included the CTL marker CD8 and the Treg marker FOXP3 (Fig. 1A). Besides cells expressing a single marker (i.e., CD8+, CTLs and FOXP3+, Treg cells), co-expression of CD8+ and FOXP3+ was also detected. These CD8+FOXP3+ double positive cells have been described as Treg cells with phenotypic resemblance to classical CD4+FOXP3+ Treg cells (18,19), however, for the purposes of this paper they have been analyzed separately. Infiltration of T cells was heterogeneous within each tumor type, with no specific type-dependent differences (Fig. 1B-E). The exception was CTLs, which were detected in roughly half of ESS and UUS cases (Fig. 1F and 1G) but were completely absent in all other cases.

**Figure 1.**
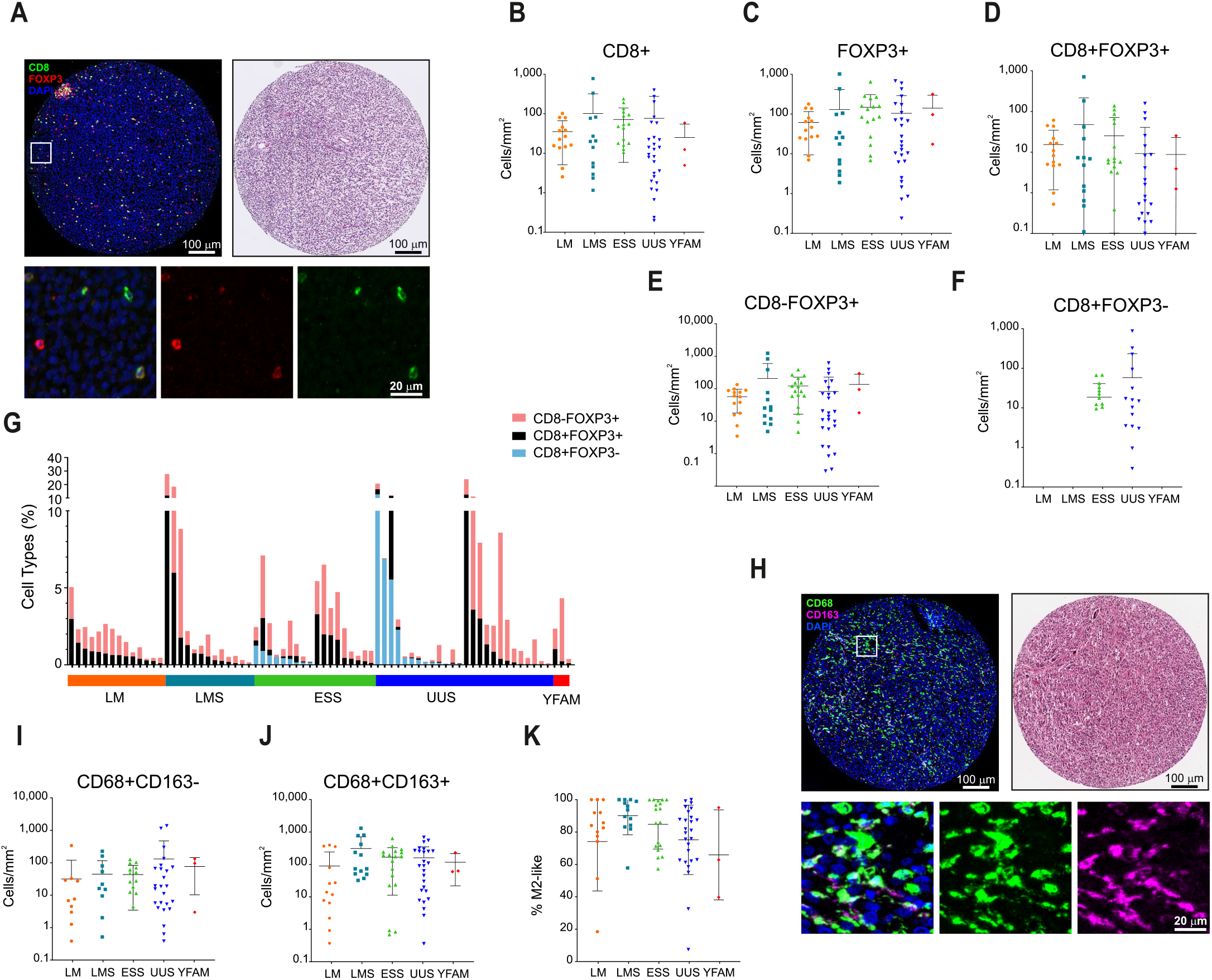
T cell and macrophage infiltration is highly heterogeneous in uterine mesenchymal tumors. **A**, IF staining of a representative example of a tumor with infiltrated T cells (left), and its corresponding H&E staining (right), and representative images of single and double staining for CD8 and FOXP3 (bottom). **B-F**, Quantification of T cell densities shows heterogeneity in overall CD8+ (**B**), overall FOXP3+ (**C**), CD8+FOXP3+ (**D**), and CD8−FOXP3+ (**E**) in leiomyoma (LM), leiomyosarcoma (LMS), endometrial stromal sarcoma (ESS), undifferentiated uterine sarcoma (UUS), and YFAM translocation-bearing ESS. **F**, CD8+FOXP3− cells are found in ESS and UUS but not in LM or LMS tumors. **G**, Proportion of T cell types in individual patients. **H**, Representative IF staining of macrophages (left), their corresponding H&E staining (right), and representative examples of single and double CD68- and CD163-expressing cells (bottom). **I-J**, Quantification of macrophage densities shows heterogeneity in overall CD68+CD163− (**I**) and CD68+CD163+ (**J**) infiltration. **K**, Percentage of macrophages with M2-like phenotype. Significance is indicated as *(P< 0.05),** (P<0.01).

Macrophages were characterized using the CD68 and CD163 markers, allowing quantification of M1-like (CD68+CD163−) and M2-like (CD68+CD163+) populations (Fig. 1H). Macrophage infiltration was heterogeneous within each tumor type, with no specific type-dependent differences (Fig. 1I and J). Yet, anti-inflammatory M2-like macrophages were more prevalent than M1-like macrophages in all tumor types, with most tumors presenting over 60% M2-like macrophages (Fig. 1K).

The expression of potentially targetable IRPs, including PD-1, PD-L1, B7-H4 and IDO1, were also characterized by immunofluorescence (Fig. 2A). Expression of PD-1, B7-H4 and IDO1 was heterogeneous in the different tumor types (Fig. 2B-D) with LMS tumors presenting a higher B7-H4 expression than ESS and UUS. Expression of PD-L1 was not detected in any group (Fig. 2E, simultaneously stained control tissue showed expected positive staining, Supplementary Fig. S1A). Further classification of cells co-expressing PD-1 with CD8 and FOXP3, indicative of T cell exhaustion, revealed that PD-1 is expressed in a subset of Treg but not in CTLs, indicating that CTLs are not exhausted in these tumors (Fig 2F, simultaneously stained control tissue showed CD8+PD-1+ expressing cells, see Supplementary Fig. S1B). Moreover, the proportion of exhausted Treg was generally low (Fig. 2G; 7.0% in CD8+FOXP8+ and 3.9% in CD8−FOXP3+). Interestingly, a large proportion of PD-1+ cells did not express either CD8 or FOXP3. As CD8−FOXP3+ and CD8-FOXP3-PD-1+ cells are likely to be CD4+ Treg and exhausted CD4+ T cells, respectively, we assessed the CD4+ cell infiltration in these tumors by immunohistochemistry (Fig. 2H). Infiltration of CD4+ cells was heterogeneous in all tumor types and positively correlated with CD8-FOXP3+ and CD8-FOXP3-PD-1+ cell infiltration (Fig. 2I, J), suggesting that a considerable proportion of these cells are CD4+.

**Figure 2.**
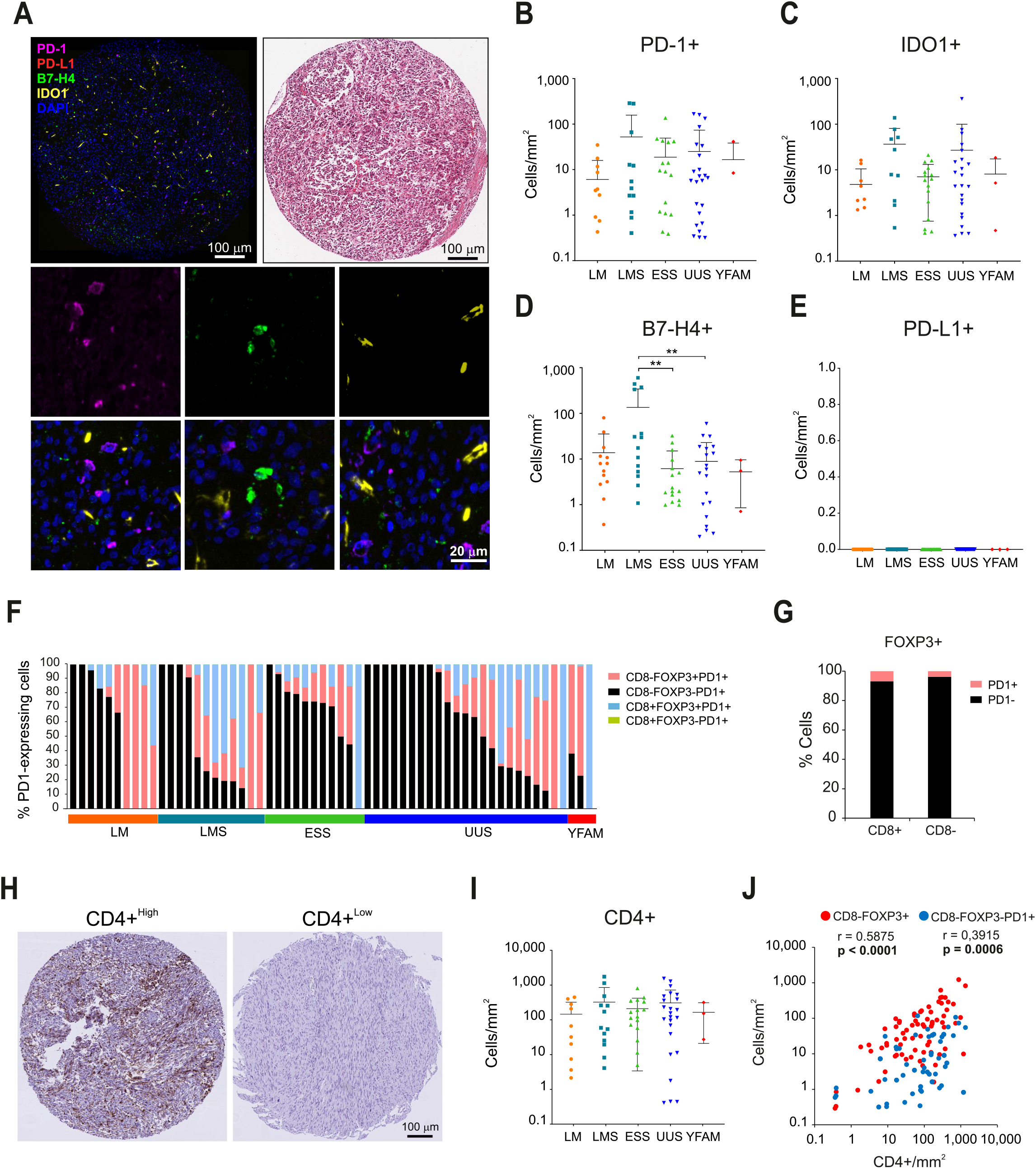
The expression of immune regulatory proteins (IRPs) is heterogeneous in uterine mesenchymal tumors. **A**, Representative IF staining of an IRPs-expressing tumor (left), its corresponding H&E staining (right) and representative images of cells expressing PD-1, B7-H4, and IDO1 (bottom). **B-E,** Quantification of IRP-expressing cells shows heterogeneous expression of PD-1+ (**B**), IDO1+ (**C**), B7-H4+ (**D**), whereas PD-L1 is not detected in any tumor group (**E**). **F,** Percentage types of PD-1 expressing cells. **G,** PD-1 expression in FOXP3+ cells. **H,** Representative examples of IHC staining for CD4+ T cells representing high and low infiltration tumors. **I**, Quantification of CD4+ T cell density shows heterogeneity in CD4+ expression in all tumor types. **J**, Scatter plot shows the significant correlation between IHC stained CD4+ and IF stained CD8-FOXP3+ and CD8-FOXP3-PD1+ T cells. Significance is indicated as *(P< 0.05), ** (P<0.01).

### Uterine mesenchymal tumors can be divided into four intrinsic immune cell subtypes

To better understand the nature of the immune cell microenvironment, we performed an unsupervised clustering of individual patients based on immune cell infiltration and expression of IRPs. This analysis resulted in clustering of patients based on the tumor immune cell infiltrate rather than the specific tumor type (Fig. 3A). Three distinct immune cell groups were identified. These immune groups were defined by Macrophages, Treg, and IRP together with CTLs (Fig. 3B). Patient clustering revealed two major clusters. One cluster included tumors with high immune cell infiltration (hot tumors; 4 LMS, 3 ESS, 6 UUS, and 3 LM). A second major cluster generally presented low immune cell infiltrates (cold tumors; 9 LMS, 13 ESS, 20 UUS, 3 YFAM). Clustering of cold tumors further revealed three specific subclusters. These included a Treg-high (immune excluded; 1 LMS, 4 ESS, 1 UUS, 1 YFAM, and 2 LM), a group of tumors expressing IRPs (immune ignored; 4 LMS, 4 ESS, 11 UUS, and 8 LM) and a third subgroup expressing either exhausted Treg or some IRP/CTLs markers (non-specific; 4 LMS, 5 ESS, 8 UUS, 2 YFAM, and 1 LM). These results again indicate that immune cell infiltrates are largely independent of the current tumor subtype classification.

**Figure 3.**
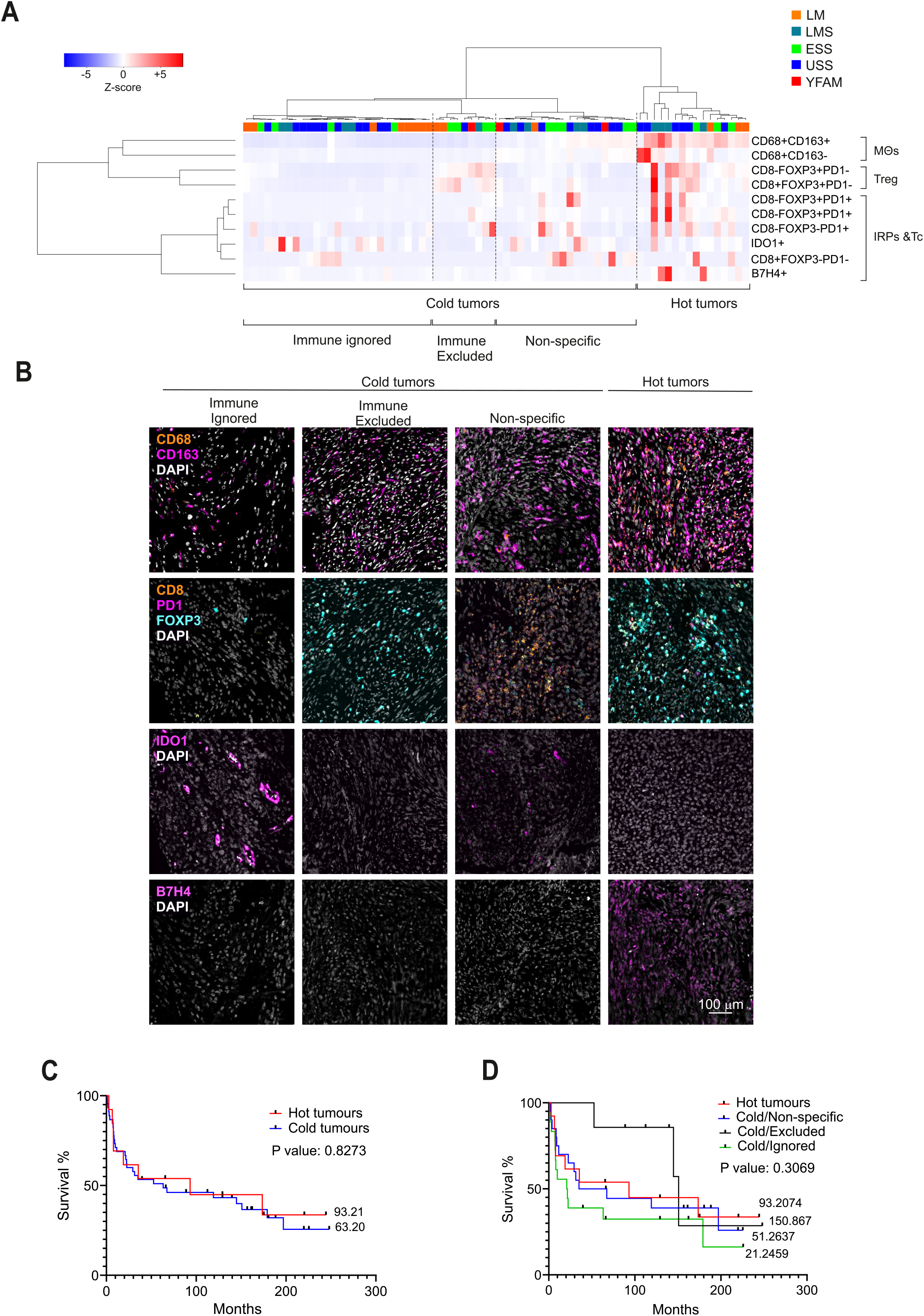
Unsupervised hierarchical clustering of immune marker expression reveals 4 distinct immune groups independently of tumor types. **A**, unsupervised hierarchical clustering of mesenchymal uterine tumors based on expression of immune markers. **B**, IF images of tumors representing the 4 immune groups represented in the unsupervised hierarchical clustering. **C**, Kaplan-Meier curves of patients with hot and cold tumors defined in (**A**). **D**, Kaplan-Meier curves of patients grouped based on the 4 distinct immune groups defined in (**A**).

Finally, we assessed the prognostic value of these patient groups, excluding the benign cases. Comparing patient overall survival between hot and cold tumors did not show any major differences in overall patient survival (Fig 3C). Similarly, overall survival (OS) did not show statistical differences when cold tumors were further divided into the three subgroups (Fig. 3D).

### Correlation with clinicopathologic data shows that Treg and M1-like macrophage infiltration are associated with improved overall survival

We next evaluated the relationship between OS and individual immune cell infiltrates. Considering all cases, improved OS was significantly associated with the presence of FOXP3+ Treg cells (Fig. 4A-C) but not with CD8+ or CD4+ cells (Fig. 4D and E and Supplementary Fig. S2A). Moreover, abundance of M1-like, but not M2-like, macrophages was associated with better OS (Fig. 4F and Supplementary Fig. S2B). Dividing the cases by tumor subtype, the Treg survival improvement was observed in UUS, and as a non-significant trend in ESS (p=0.0269 and p=0.0954, respectively), but not in LMS (Supplementary Fig. S3). The M1-like macrophage prevalence was associated with survival improvement, specifically in UUS patients (p=0.0296, Supplementary Table 3). Neither PD-1, IDO1, nor B7-H4 expression showed general or tumor type-specific prognostic value (Supplementary Fig. S4A-C and Supplementary Table 4). However, absence of PD-1-expression in CD8+ and CD8-Tregs was a marker of good prognosis (Fig. 4G-J). Contrarily, the infiltration of PD1+ cells that do not express either CD8 or FOXP3 was associated with improved OS (Fig. 4K).

**Figure 4.**
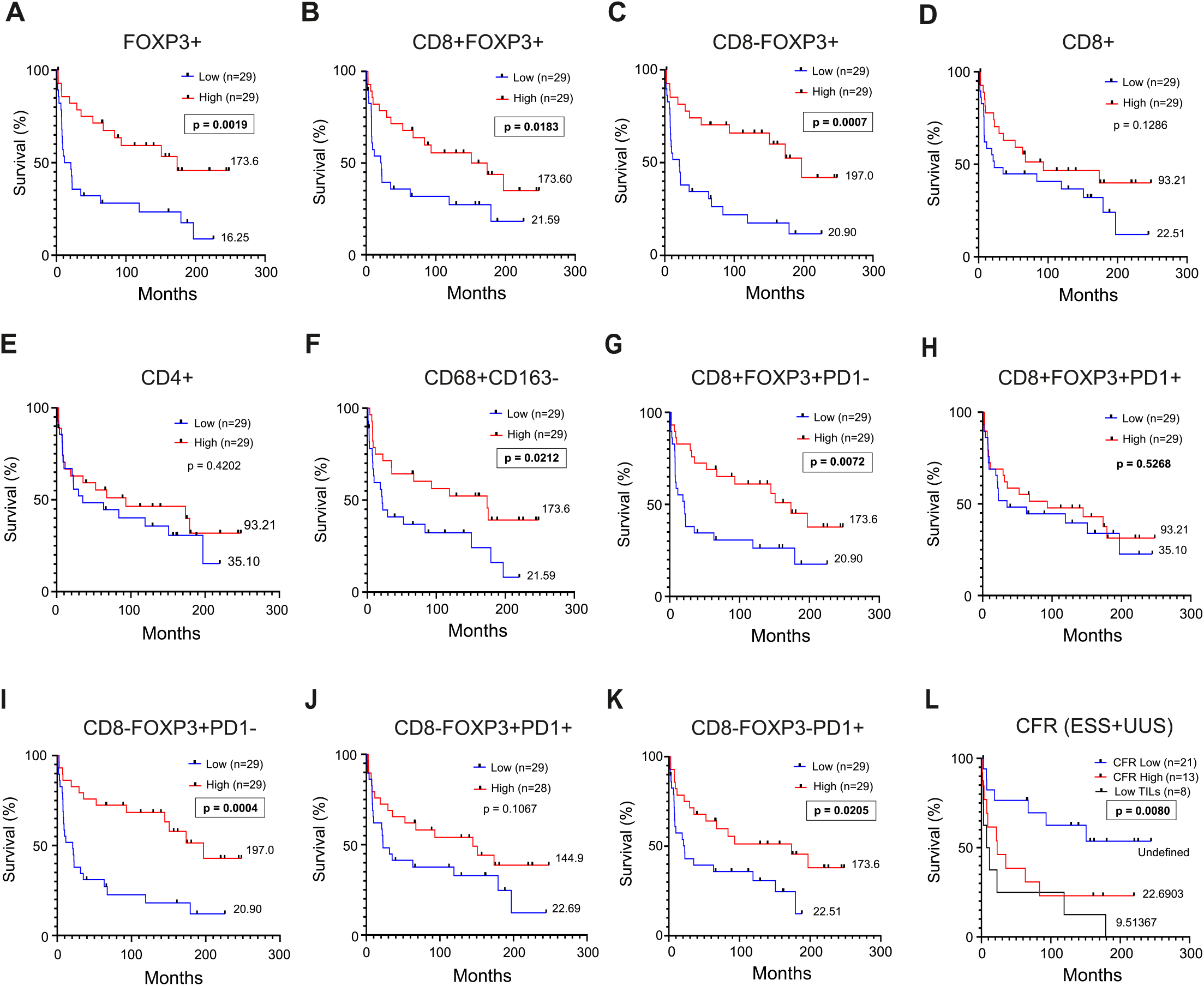
Treg cell and M1-like macrophage density is associated with better survival. **A-K,** Kaplan-Meier curves indicating patient overall survival. Patients were grouped based on median density of specific cell subtypes regardless of diagnosis, excluding benign leiomyomas. Median survival is indicated for each patient group. **A-E**, Kaplan-Meier curves of patients grouped based on distinct T cell markers indicate FOXP3 as a marker of good prognosis. **F**, Kaplan-Meier curves based on M1-like type macrophage density show CD68+CD163− cells to be a marker of improved survival. **G-J** Kaplan-Meier curves based on distinct Treg cell populations classified according to their PD-1 expression, show that only PD-1-negative Treg cell infiltration is associated with good prognosis. **K** Kaplan-Meier curves indicating overall survival based on density of CD8-FOXP3-PD1+ cells. **L,** Kaplan-Meier curves showing overall survival based on CD8+FOXP3-/FOXP3+ ratio (CFR) including ESS and UUS cases (Low TILs group corresponds to CD8+FOXP3- and FOXP3+ density below the 40^th^ percentile).

In other tumor types, the CD8+/FOXP3+ ratio (CFR) has been shown to have a stronger association with patient survival than individual cell types (20–22). We investigated the impact of this ratio on survival in ESS and UUS as these contained both CTLs and Treg cells. Cases with few tumor infiltrating lymphocytes (Low TILs) were considered as a third group. Low CFR (< 1) (i.e., tumors with a larger proportion of Treg than CTLs cells) was associated with significantly improved survival, compared to both CFR^High^ and low TILs (Fig. 4L). This association was stronger in ESS than in UUS (Supplementary Fig. S5A and B). Together, these data show that infiltration of Tregs (either as an absolute density or CFR), M1-like macrophages, and CD8-FOXP3-PD1+ (i.e., presumed CD4+PD1+ cells), are associated with improved overall survival, suggesting that these cells are involved in modulating tumor aggressiveness.

### High CFR associates with ECM signaling genes

To further understand the biological underpinnings of the observed survival differences, we first compared the gene expression of UUS (n=22) between CFR^High^ and all other tumors, which revealed 267 differentially expressed genes. Specifically comparing CFR^High^ and CFR^Low^ tumors resulted in 524 differentially expressed genes, indicating that these tumors differ considerably at the gene expression level. Contrarily, transcriptomic analysis of tumors expressing high and low M1-like macrophages showed 18 differentially expressed genes, indicating that these tumor groups are transcriptionally similar (Supplementary Table 5). Subsequently, we conducted a pathway analysis to uncover major pathway alterations between CFR^High^ and other tumors. Major alterations were found in pathways that included extracellular matrix organization, proteoglycans in cancer, regulation of cell adhesion, and integrin-mediated signaling (Fig. 5A). Similarly, independently comparing tumors with high vs low CTLs or tumors with high vs low Treg cells, showed ECM-related pathways as the most significantly altered ones (Supplementary Fig. S6). MCODE analysis for protein-protein interactions showed two densely connected neighborhoods (Fig. 5B). Gene ontology enrichment analysis of these two MCODE neighborhoods linked MCODE1 with non-integrin membrane-ECM interactions, integrin pathway, and laminin interactions, and MCODE2 with smooth muscle contraction, muscle contraction, and actin cytoskeleton organization. These results indicate that differences in tumor CFR are related to ECM-cell adhesion and signaling at the transcriptional level.

**Figure 5.**
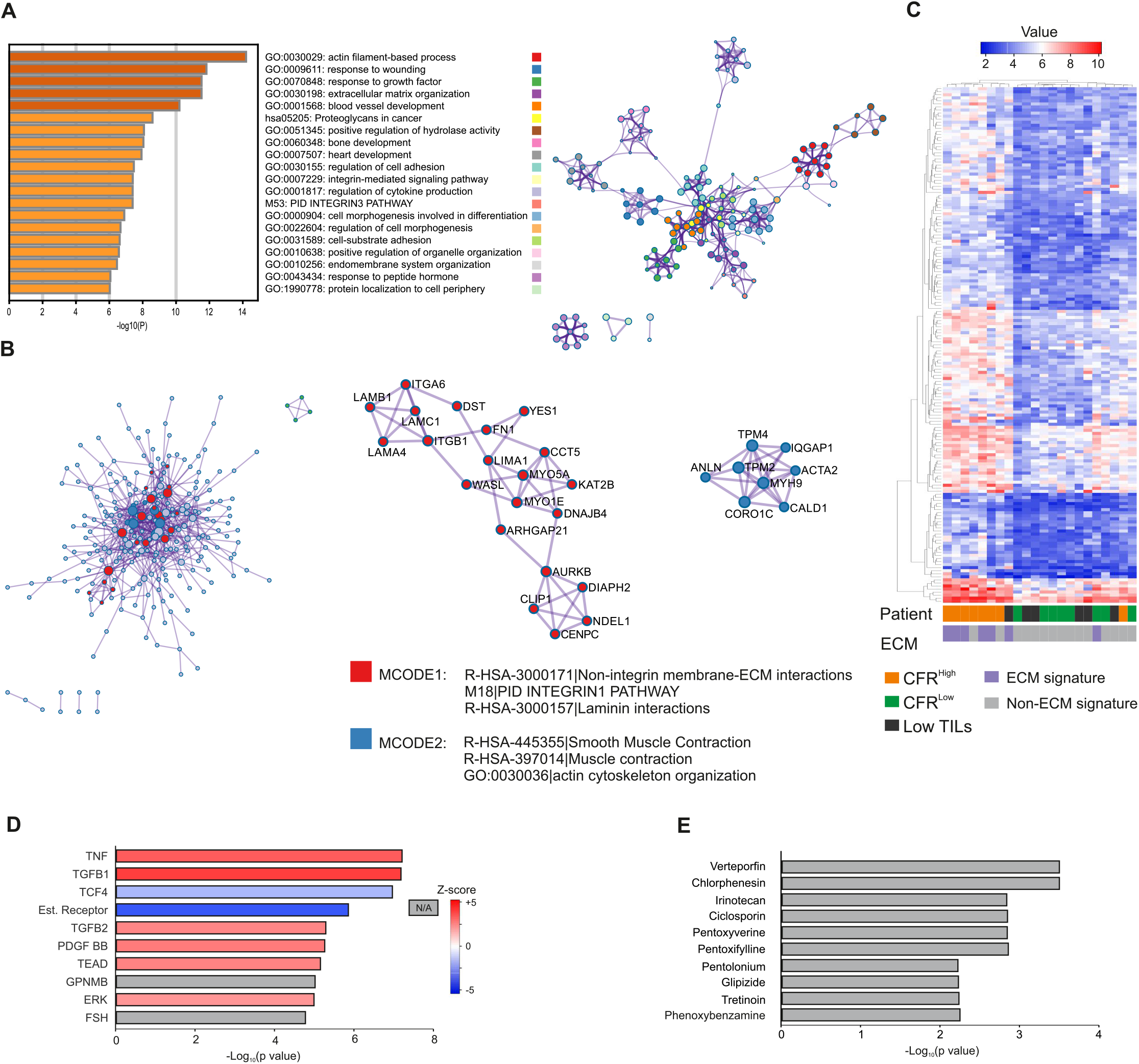
Tumors with high CD8+FOXP3-/FOXP3+ ratio show upregulation of ECM-Integrin interaction pathways in our UUS cohort. **A**, Metascape pathway analysis reveals the most significantly altered pathways in the CFR^High^ group (CD8+FOXP3-/FOXP3+ > 1) and their interactions. **B**, MCODE-identified neighborhoods of densely connected proteins based on upregulated genes in the high CFR group showing their corresponding enriched Gene Ontologies. **C**, Heat map of gene expression of significantly altered genes in the CFR-defined groups with patient clustering indicating their corresponding ECM signature as defined by Binzer-Panchal et. al. **D**, Ingenuity pathway analysis (IPA) indicates the potential upstream regulators of the differential gene expression between CFR high and low tumors. **E**, Connectivity Map analysis shows potential gene expression-altering drugs for the upregulated genes in the CFR^High^ compared to CFR^Low^ groups.

Previously, we identified an ECM gene signature in a subgroup of aggressive UUS tumors (15). Comparing tumor CFR and ECM signature revealed that 5 out of 8 CFR^High^ tumors presented the ECM gene signature (Fig. 5C). Finally, to further uncover the biological characteristics of CFR^High^ tumors, we identified likely upstream regulators and drugs targeting the genes upregulated in these tumors by Ingenuity Pathway Analysis and Connectivity Map, respectively. These analyses revealed TNF, TGFB1/2, PDGF-BB, and TEAD as upstream regulators (Fig. 4D), all of which are involved in tissue fibrosis (23–26). Furthermore, verteporfin and chlorphenesin were uncovered as potential gene perturbing drugs (Fig. 4E). Interestingly, verteporfin is a suppressor of YAP-TEAD complex downstream of ECM-integrin signaling, and chlorphenesin is a muscle relaxant (27,28). Altogether, these results suggest ECM-signaling as a regulator of T cell populations in uterine sarcomas.

### CFR^Low^ tumors show reduced ECM protein expression and YAP nuclear localization

To validate the ECM gene expression differences, we analyzed the protein expression of collagen I, VI, fibronectin, and MMP14, as these proteins are significantly enhanced in tumors with the ECM gene signature. In UUS tumors, the expression of collagen I, collagen VI, and fibronectin was lower in CFR^Low^ than in the CFR^High^ group, although fibronectin was not significant (p = 0.0699) (Fig. 5A). Similarly, the expression of these proteins was lower in CFR^Low^ ESS tumors compared to CFR^High^, although MMP14 expression similar (Fig. 6B). To validate the possible implication of YAP signaling in defining the tumor CFR, we compared the YAP nuclear to cytoplasmic ratio in the different CFR-based groups. In conjunction with ECM protein expression, the proportion of nuclear YAP, suggesting YAP activation, was higher in CFR^High^ compared to CFR^Low^ and Low TILs, significantly in UUS tumors (Fig. 6C and D). Together, these data show a correlation between ECM expression, YAP nuclear localization, and high CTLs/low Treg infiltration in ESS and UUS, suggesting a mutual regulation of these components in the tumor microenvironment.

**Figure 6.**
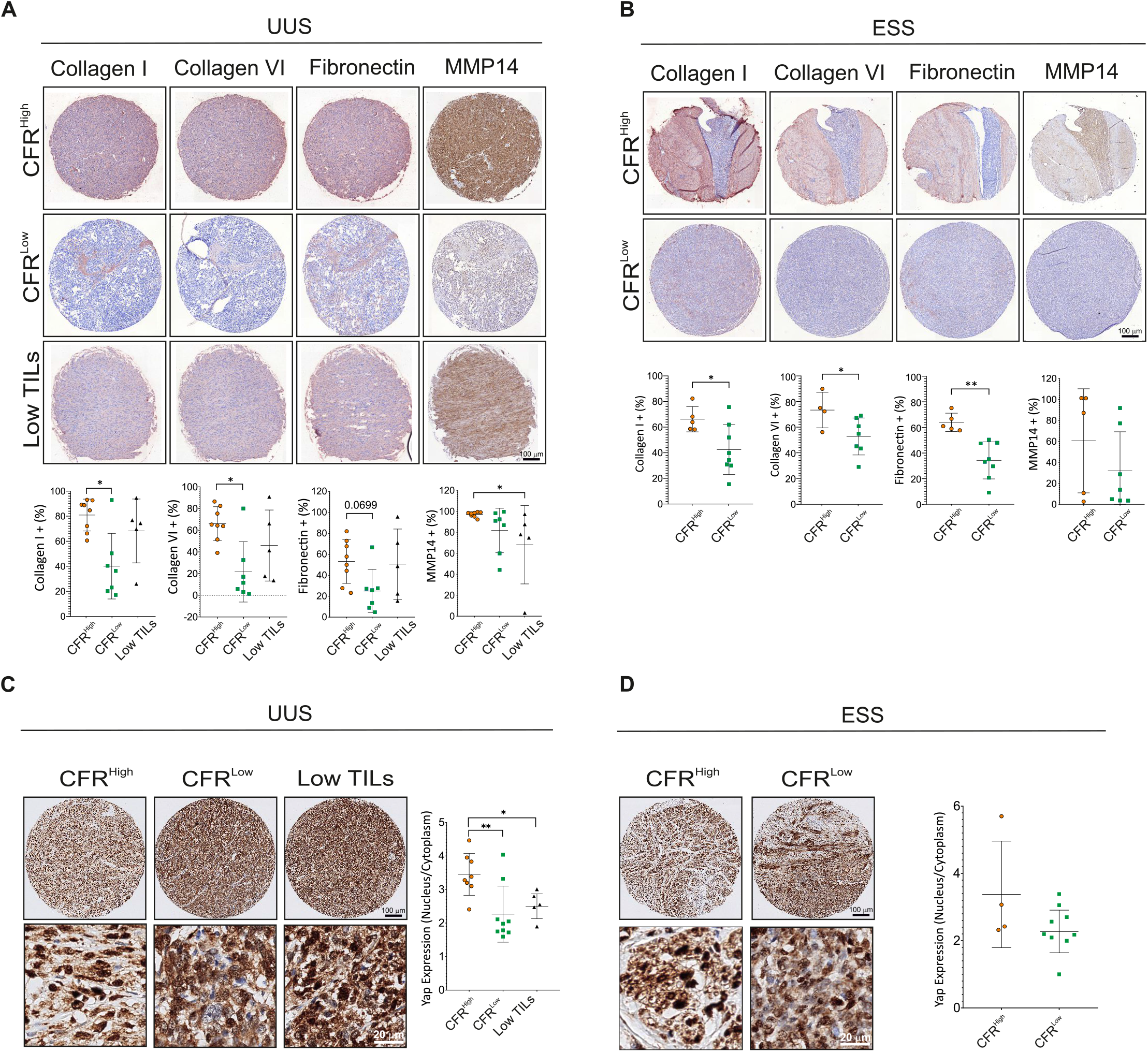
ECM protein and nuclear YAP expression is enhanced in UUS and ESS tumors with high CFR. **A**, **B**, Representative IHC images and quantification of indicated protein expressions show higher expression in the CFR^High^ group in UUS (**A**) and in ESS patients (**B**). **C,D,** Representative IHC images of YAP expression and quantification of its nuclear/cytoplasmic ratio shows higher ratio in the CFR^High^ patient group in UUS (**A**) and in ESS patients (**B**). Significance test is indicated *(P< 0.05), ** (P<0.01).

## Discussion

Uterine sarcomas are aggressive and rare mesenchymal tumors that currently lack effective treatment. The biology of sarcomas and their interaction with their microenvironment is complex and influences the potential utility of immunotherapy. There is a critical need to understand the sarcoma immune microenvironment and the molecular mechanisms behind it, in order to identify new therapies.

In the work reported here, we have used multiplex immunofluorescence to identify TIL and macrophage populations across a variety of uterine sarcomas, and to investigate the expression of potentially targetable IRPs. Further, we have used transcriptomic and protein level data to identify potential pathways involved in the observed TIME differences. These experiments have led to several surprising findings.

First, the immune microenvironment is largely independent of tumor type. Previous studies have shown that immune cell accumulation is associated with diverse sarcoma subtypes, where dominant infiltration of TAMs was observed in UUS and undifferentiated pleomorphic sarcomas (29,30). In line with these reports, our results indicate that the immune microenvironment depends on factors other than the tumor type. This is seen when considering the infiltration of individual immune cell types and seen again in our identification of intrinsic immune cell subgroups. This finding carries an important translational relevance; namely, that immunotherapy trials should be performed with a well-planned translational component, so that observed responses can be related to the TIME in those tumors. These hypothetical predictive biomarkers may then be “agnostic” to sarcoma subtype and should be evaluated as such. This could greatly assist with the clinical management of sarcomas, where the large number of histopathologic subtypes can make traditional “subtype specific” trials difficult. Instead, trials should focus on the varieties of TIME and intrinsic immune cell subgroup present.

Second, we have demonstrated that the nature of the immune cell infiltrate has an influence on overall survival. The presence of non-exhausted Treg cells and M1-like macrophages was associated with better prognosis. This association was observed in both classical CD8- and in CD8+ Treg cells. Classifying tumors according to their CD8 to FOXP3 ratio (CFR) indicated that low CFR is associated with improved survival. This result is counterintuitive, given that in most tumors it is the presence of CTLs that is associated with a better prognosis. Our results indicate that cytotoxic T cell infiltration plays a much less important role in uterine sarcomas. The current knowledge of Treg cell function in uterine sarcomas is limited. However, our data indicate that these cells play a central role in uterine sarcoma progression and are associated with better prognosis. Contrarily, a previous study identified Treg cells to associate with high tumor grade and poor survival in a cohort including various soft tissue sarcoma types (31), suggesting that our findings are limited to uterine sarcomas. The good prognosis associated with Treg cells opposes the general tumor-promoting role of these cells, but similar observations have been made in colorectal cancer, head and neck carcinoma, and Hodgkin lymphoma, among others (32). One possible explanation for these effects is that a significant proportion of the FOXP3+ cells detected are not suppressive Treg cells, rather non-suppressive cells with low expression of FOXP3, which secrete pro-inflammatory cytokines, and are associated with better prognosis in colorectal cancer (33). As these non-Treg FOXP3+ cells lack the expression of CD45RA, including this marker for further characterization of this cell population may shed light on the function of FOXP3+ cells in uterine sarcomas and other tumors.

Third, the results from our transcriptomics analysis indicate that pathways such as extracellular matrix organization, proteoglycans in cancer, regulation of cell adhesion, and integrin-mediated signaling are associated with the observed differences in TIME. Predicted drug regulators of these altered genes highlighted verteporfin and chlorphenesin. Verteporfin is an inhibitor of the YAP-TEAD transcription factor complex (34). There is currently a clinical trial planned for the use of liposomal verteporfin in the treatment of recurrent glioblastoma multiforme, and previous reports indicate that verteporfin can modulate the TIME through PD-L1 inhibition (35). Although our findings show that uterine sarcomas present low PD-L1 expression, independent responses of the YAP-TEAD inhibition may lead to the regulation of the TIME in these tumors.

Finally, analysis of protein expression confirms the role of the ECM regulatory proteins in the observed differences in immune cell infiltration. Furthermore, the proportion of nuclear vs. cytoplasmic YAP indicates that YAP signaling may play a role in this process. YAP is clearly a central regulator of ECM signaling and is involved in sarcomagenesis (36,37). Thus, our data suggests that YAP is involved in uterine sarcoma progression and may have immune regulatory functions. Moreover, these results show that, independently of tumor type, tumors with an ECM signature present similar immune microenvironment. Thus, characterizing tumors for specific TME signatures might be a better approach to precision immunotherapy than solely using traditional histopathologic subtypes.

These results have numerous translational implications. First, they indicate that clinical trials evaluating immune therapies will need to include a strong translational component to identify what immune cells underlie any potential observed response. Furthermore, any immune cell signature that can be related to a response should then be evaluated in a tumor agnostic manner, given the heterogeneity observed. The central role of YAP and ECM pathways in these differences indicate that therapies focusing on modulation of the ECM should be considered.

## Authors’ Contributions

### Conception and design

J. W. Carlson

### Development of methodology

J. W. Carlson

### Acquisition of data (provided animals, acquired and managed patients, provided facilities, etc.)

E. Hardell, G. Kokaraki, L. Moyano-Galceran, N. Mitsios, J. Mulder, A. Isaksson, K. Lehti, J.

W. Carlson

### Analysis and interpretation of data (e.g., statistical analysis, biostatistics, computational analysis)

O. Gultekin, J. Gonzalez-Molina, E. Hardell, D. Sarhan, K. Lehti, J. W. Carlson

### Writing, review, and/or revision of the manuscript

O. Gultekin, J. Gonzalez-Molina, L. Moyano-Galceran, N. Mitsios, D. Sarhan, K. Lehti, J. W. Carlson

### Administrative, technical, or material support (i.e., reporting or organizing data, constructing databases)

O. Gultekin, J. Gonzalez-Molina, E. Hardell, J. Mulder, A. Isaksson, J. W. Carlson

### Study supervision

K. Lehti, J. W. Carlson

## Acknowledgments

J. W. Carlson is supported by The Swedish Cancer Society and The Radiumhemmets Forskningsfonder J. Gonzalez-Molina is supported by Barncancerfonden D. Sarhan is supported by The Swedish Cancer Society and KI foundations

