## Supplementary material for "Regulatory FOXP3+ T cells in uterine sarcomas are associated with favorable prognosis, low extracellular matrix expression and reduced YAP activation": Supplemetary Figure legends

**Supplementary Table 1**

Summary of the details of the antibodies used for immunofluorescence and immunohistochemistry. Human protein atlas is represented as HPA.

**Supplementary Table 2**

Clinical Data of Cohort including age at diagnosis, disease stage at diagnosis, and survival.

**Supplementary Table 3**

Summary of median survival and P values for patients grouped according to their M1-like or M2-like macrophage infiltration in ESS, UUS, and LMS tumors.

**Supplementary Table 4**

Summary of median survival and P values for patient grouped according to their immune regulatory protein expression in ESS, UUS, and LMS tumors.

**Supplementary Table 5**

List of differentially expressed genes (> 2-fold) between UUS tumors with high and low infiltration of M1-like macrophages.

**Supplementary Figure S1**

PD-L1 expression and CD8+PD1+ cells were detected in control endometrial carcinoma tissues but not in uterine sarcoma. **A**, Example images of PD-L1 staining and QuPath-based cell type detection (bottom right) showing PD-L1+ cells in endometrial carcinoma tissue. **B**, Example images of enometrial carcinoma tissue presenting CD8+PD1+ cells. Arrowhead indicates an example of a CD8+PD1+ cell.

**Supplementary Figure S2**

Kaplan-Meier curves showing infiltration of CTLs (CD8+FOXP3-) and M2-like macrophages (CD68+CD163-), which are not prognostic in uterine sarcomas.

**Supplementary Fıgure S3**

Treg cell density is associated with better survival in UUS and ESS. **A-C**, Kaplan-Meier curves showing overall survival for the indicated Treg cell subgroups in UUS, ESS, and LMS patients. Median Survival is indicated for each group.

**Supplementary Figure S4**

Kaplan-Meier curves of patient groups based on expression of immune regulatory proteins PD-1 (**A**), IDO1 (**B**), and B7-H4 (**C**) showing that the general expression of these proteins does not have prognostic value in uterine sarcomas. Median Survival is indicated for each group.

**Supplementary Figure S5**

High CD8+FOXP3-/FOXP3+ ratio (CFR) is associated with poor survival. Kaplan-Meier curves of ESS (A) and UUS (B) patients grouped based on their CFR. UUS tumours include a low-TIL group defined as tumors with T cell infiltration below the 40th percentile of all tumors analyzed. Median Survival is indicated for each group.

**Supplementary Figure S6**

Tumors with distinct cytotoxic T cell and regulatory T cell infiltration show differential expression of extracellular matrix-related genes. **A**, **B** Metascape pathway analysis reveals the most significantly altered pathways between tumors with high and low cytotoxic T cell infiltration (**A**), and regulatory T cell infiltration (**B**).
