## Supplementary Figures for "Regulatory FOXP3+ T cells in uterine sarcomas are associated with favorable prognosis, low extracellular matrix expression and reduced YAP activation"

Supplementary Table 1

| Antibody | Company | Product No. | Dilution | Technique |
|---|---|---|---|---|
| CD8 | Dako | M 7103 | 1 : 100 | IF |
| FOXP3 | EuroMAbNET | 236A/E7 | none | IF |
| PDCD1 / CD279 (PD1) | HPA | HPA035981 | 1 : 250 | IF |
| CD68 | Dako | M 0876 | 1 : 100 | IF |
| CD163 | HPA | HPA046404 | 1 : 1600 | IF |
| PDL1 | Dako | M 3653 | 1 : 100 | IF |
| IDO1 | HPA | HPA023149 | 1 : 50 | IF |
| VTCN1 (B7-H4) | HPA | HPA054200 | 1 : 70 | IF |
| CD4 | Ventana | 790-4423 | none | IHC |
| Collagen I | Abcam | ab34719 | 1 : 200 | IHC |
| Collagen VI | Abcam | ab6588 | 1 : 200 | IHC |
| Fibronection | Sigma | F3649 | 1 : 200 | IHC |
| MMP14 | Millipore | MAB3328 | 1 : 100 | IHC |
| YAP1 | Abcam | ab56701 | 1 : 1000 | IHC |

Supplementary Table 2

| | Leiomyosarcoma | Low grade endometrial stromal sarcoma | Undifferentiated uterine sarcoma | YWHAE-FAM22 translocated sarcoma | Leiomyoma |
|---|---|---|---|---|---|
| **Number of cases** | 13 | 16 | 26 | 3 | 13 |
| **Age at diagnosis** (mean, years) | 61.4 | 54.4 | 62.2 | 59.1 | |
| **Stage** | | | | | |
| 1 | 6 | 12 | 10 | 3 | |
| 2 | 0 | 3 | 3 | 0 | |
| 3 | 0 | 1 | 4 | 0 | |
| 4 | 4 | 0 | 2 | 0 | |
| N/A | 3 | 0 | 7 | 0 | |
| **Time to last follow up** (months) | 59.8 | 96.0 | 41.6 | 87.9 | |
| **Status at last follow up** | | | | | |
| Alive | 2 | 13 | 5 | 1 | |
| Deceased | 11 | 3 | 21 | 2 | |

Supplementary Table 3

| Tumor type | Marker | Group | n | Median survival (months) | P value (log-rank) |
|---|---|---|---|---|---|
| UUS | CD68+CD163- | High | 13 | 67.43 | **0.0296** |
| | | Low | 13 | 7.737 | |
| | CD68+CD163+ | High | 14 | 45.06 | 0.0509 |
| | | Low | 12 | 7.582 | |
| ESS | CD68+CD163- | High | 8 | Undefined | 0.9863 |
| | | Low | 8 | Undefined | |
| | CD68+CD163+ | High | 8 | Undefined | 0.7377 |
| | | Low | 8 | Undefined | |
| LMS | CD68+CD163- | High | 6 | 22.03 | 0.7196 |
| | | Low | 7 | 29.4 | |
| | CD68+CD163+ | High | 6 | 18.9833 | 0.1283 |
| | | Low | 7 | 52.5 | |

Supplementary Table 4

| Tumor type | Marker | Group | n | Median survival (months) | P value (log-rank) |
|---|---|---|---|---|---|
| UUS | IDO1 | High | 12 | 17.15 | 0.2063 |
| | | Low | 13 | 7.427 | |
| | B7H4 | High | 13 | 93.21 | 0.0606 |
| | | Low | 13 | 7.737 | |
| | PD-1 | High | 13 | 21.59 | 0.7571 |
| | | Low | 13 | 9.197 | |
| ESS | IDO1 | High | 9 | Undefined | 0.47 |
| | | Low | 7 | Undefined | |
| | B7H4 | High | 8 | Undefined | 0.6148 |
| | | Low | 8 | Undefined | |
| | PD-1 | High | 7 | Undefined | 0.4319 |
| | | Low | 9 | Undefined | |
| LMS | IDO1 | High | 5 | 173.567 | 0.3577 |
| | | Low | 5 | 9.9 | |
| | B7H4 | High | 7 | 20.9 | 0.6169 |
| | | Low | 7 | 29.4 | |
| | PD-1 | High | 6 | 44 | 0.5934 |
| | | Low | 7 | 9.9 | |

Supplementary Table 5

| ID | High Avg (log2) | Low Avg (log2) | Fold Change | P-val | FDR P-val | Gene Symbol |
|---|---|---|---|---|---|---|
| 16942270 | 4.96 | 6.39 | -2.69 | 0.0042 | 0.9998 | PTPRG |
| 16892523 | 4.75 | 6.17 | -2.68 | 0.0048 | 0.9998 | SCARNA6 |
| 16657598 | 4.03 | 5.43 | -2.63 | 0.0016 | 0.9998 | AGRN |
| 16823043 | 3.53 | 4.84 | -2.48 | 0.0067 | 0.9998 | SNHG19 |
| 16836824 | 4.47 | 5.61 | -2.21 | 0.0293 | 0.9998 | MRC2 |
| 16956386 | 4.98 | 6.08 | -2.15 | 0.0218 | 0.9998 | ROBO1 |
| 16705810 | 4.63 | 5.71 | -2.11 | 0.0393 | 0.9998 | UNC5B |
| 16668997 | 4.3 | 5.3 | -2.01 | 0.0207 | 0.9998 | OLFML3 |
| 16896836 | 3.89 | 2.88 | 2.02 | 0.0027 | 0.9998 | SLC8A1 |
| 16798951 | 5.23 | 4.17 | 2.1 | 0.0001 | 0.9998 | GREM1 |
| 17095056 | 4.98 | 3.91 | 2.1 | 0.0262 | 0.9998 | PRUNE2 |
| 16805894 | 4.3 | 3.22 | 2.12 | 0.0063 | 0.9998 | LOC646214; LOC102723432 |
| 17080788 | 6.62 | 5.38 | 2.37 | 0.0304 | 0.9998 | FBXO32 |
| 16883194 | 6.65 | 5.4 | 2.39 | 0.0348 | 0.9998 | LOC100506123 |
| 16691877 | 8.87 | 7.37 | 2.83 | 0.0093 | 0.9998 | RNU1-13P |
| 16692446 | 8.87 | 7.37 | 2.83 | 0.0093 | 0.9998 | RNU1-13P |
| 17021596 | 5.04 | 3.47 | 2.98 | 0.0261 | 0.9998 | RRAGD |
| 17016383 | 7.75 | 5.88 | 3.65 | 0.001 | 0.9998 | HIST1H4D |

Supplementary Figure S1

A

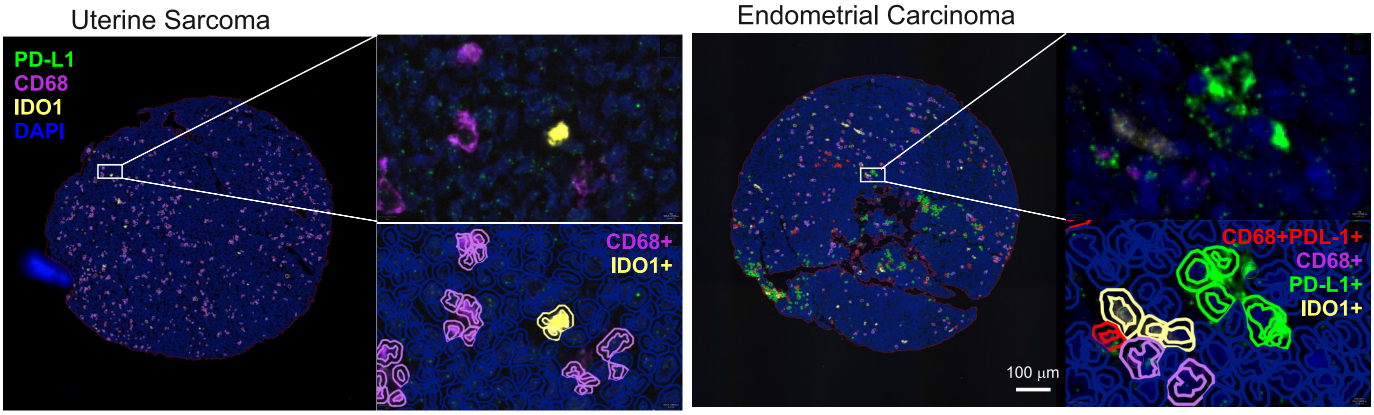

B

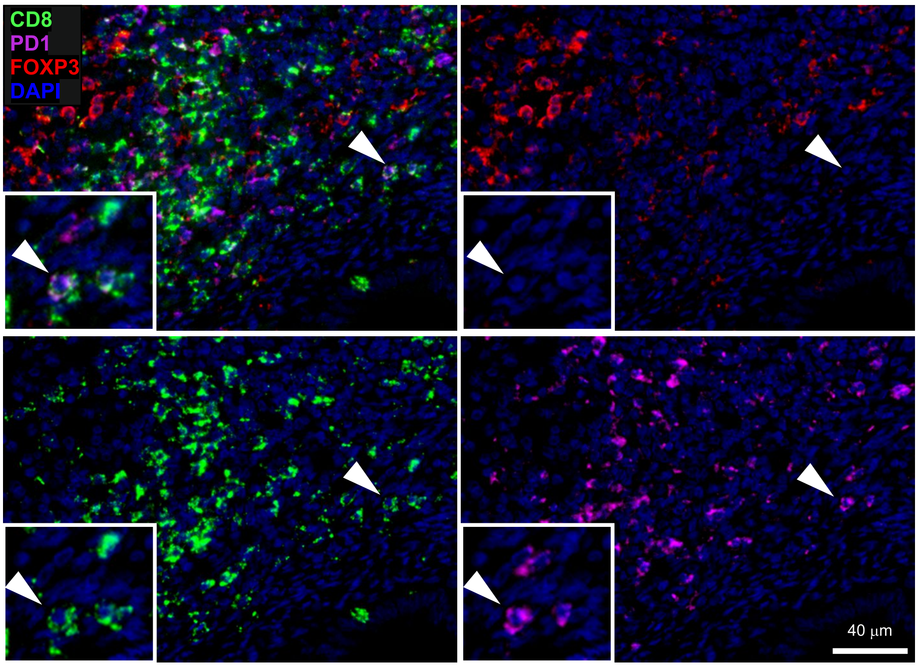

Supplementary Figure S2

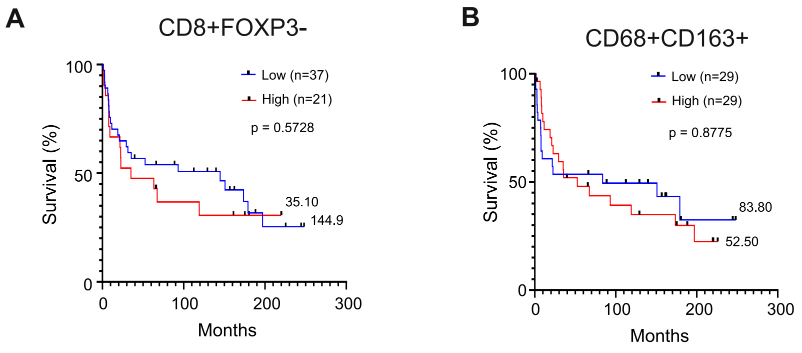

### Supplementary Figure S3

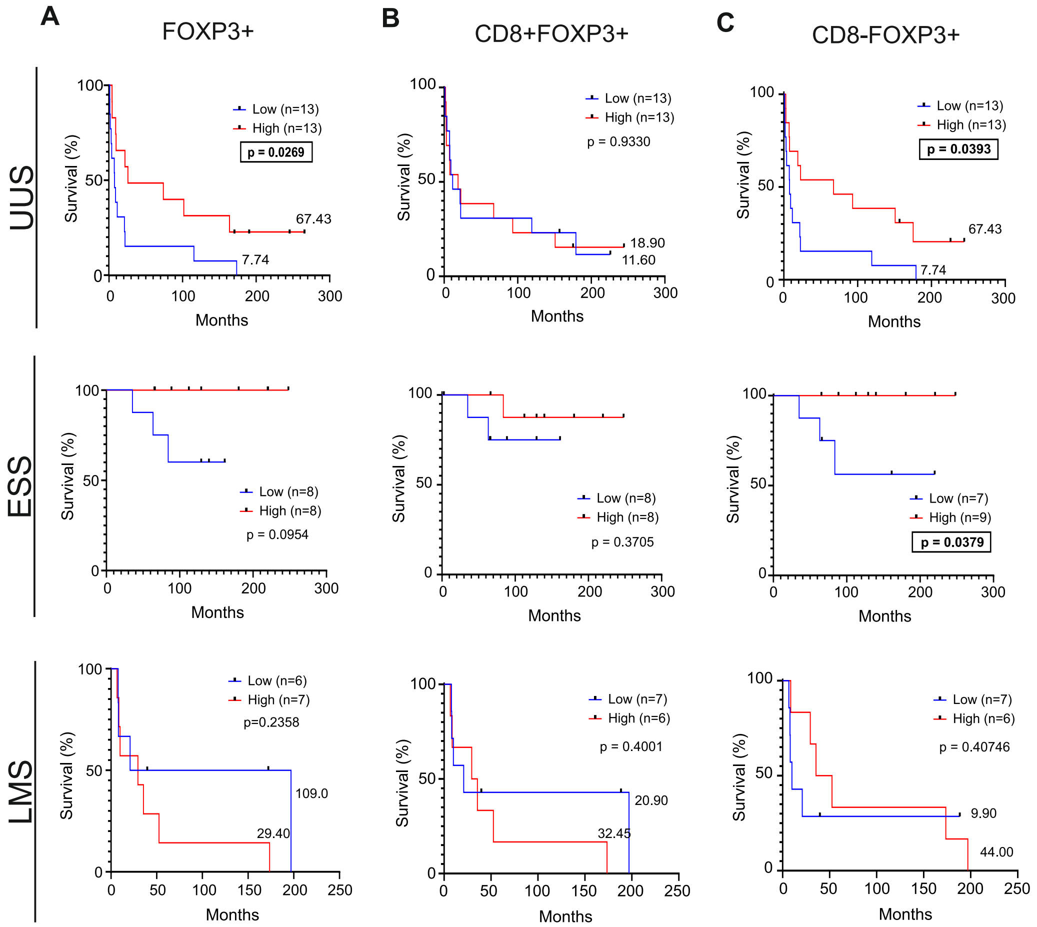

### Supplementary Figure S4

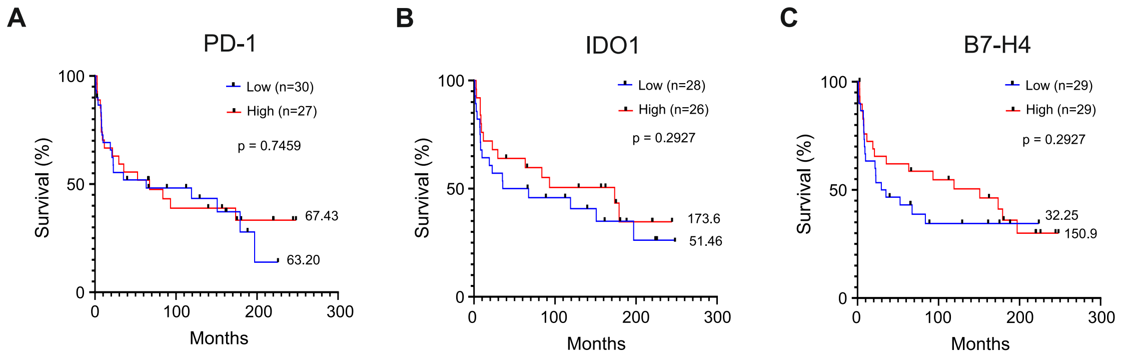

### Supplementary Figure S5

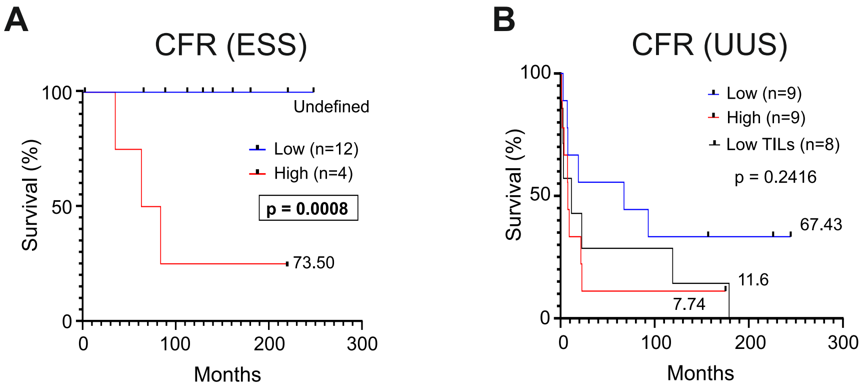

Supplementary Figure S6

A

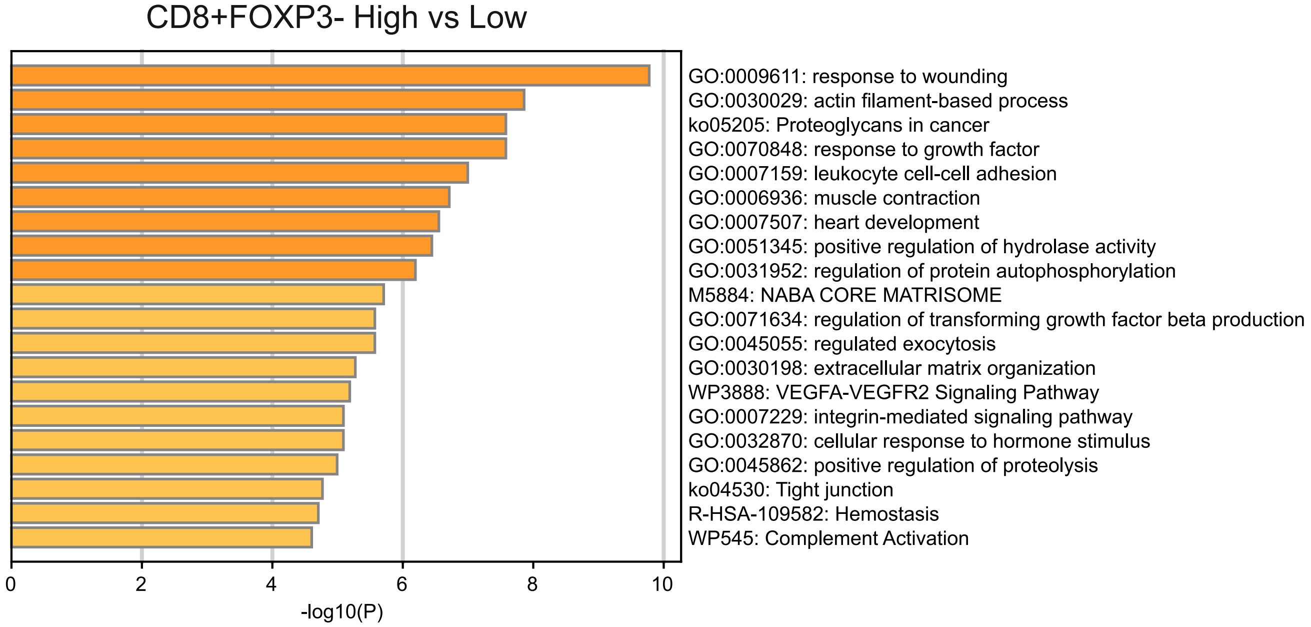

B

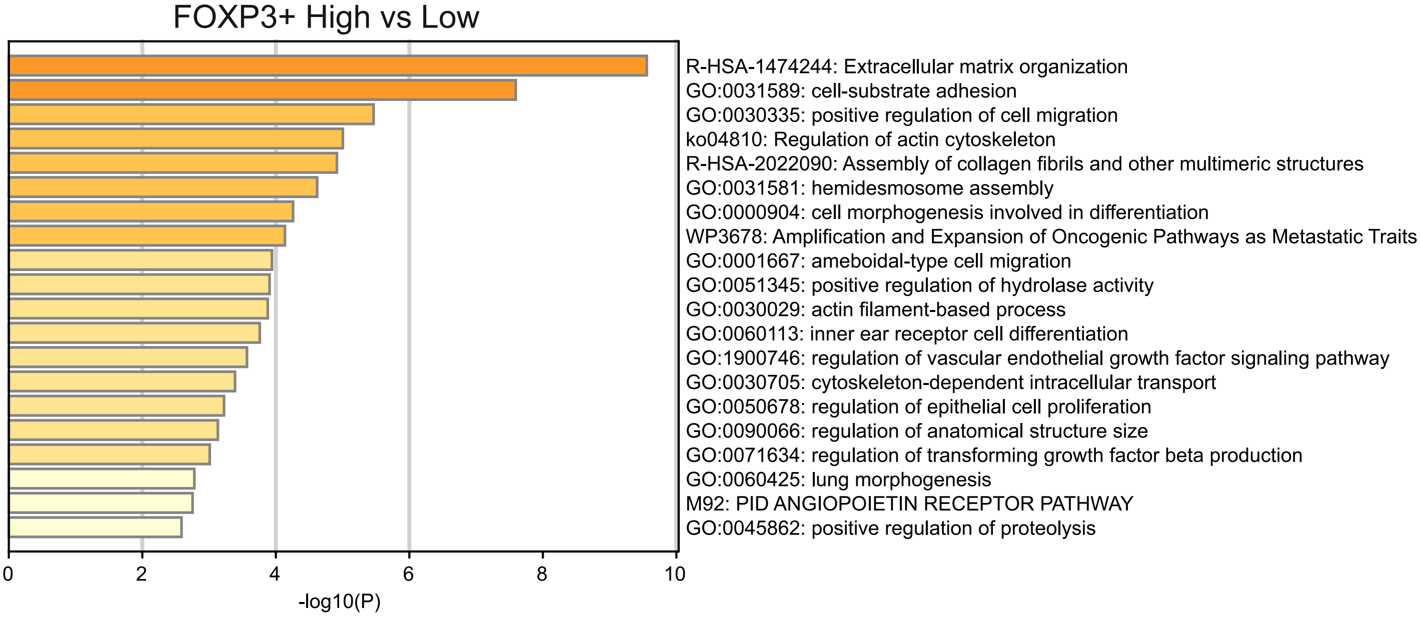
